# Spatiotemporal observation reveals metastatic tumor-driven vascular remodeling as a potential route to polyclonal colonization

**DOI:** 10.1101/2024.02.20.581134

**Authors:** Yukinori Ikeda, Hiroko Oshima, Sau Yee Kok, Masanobu Oshima, Yukiko T. Matsunaga

## Abstract

Polyclonal metastasis, which arises from clusters of circulating tumor cells, promotes metastasis development and has become a major target of metastasis inhibition. Mouse experiments have clearly verified that nonmetastatic and metastatic tumors coexist and form metastatic nests, but the detailed mechanism of extravasation remains unclear. We established a three-dimensional tumor microvessel model to investigate extravasation between nonmetastatic tumors, metastatic tumors, and mosaic tumor organoids in a mixed state by time-lapse imaging and to determine the sequential steps of the extravasation of tumor cells via vascular remodeling. This comparison revealed a new concept of extravascular invasion via vascular remodeling in metastatic carcinoma. Furthermore, the involvement of liver host cells, the hepatic stellate cells, demonstrated an interaction with metastatic cells to facilitate metastatic foci formation. Moreover, *Adam28* was highly expressed exclusively in metastatic tumor cells, suggesting its involvement in vascular remodeling. These results demonstrate the ability of metastatic tumor cells for extravasation in polyclonal metastasis, which may lead to the development of new therapeutic targets.

## 1. Introduction

Metastasis is a complex multistep process that causes the dissemination of single or clustered tumor cells to new sites. Whether metastasis is monoclonal or polyclonal has implications in influencing our understanding of metastatic potential. Polyclonal metastasis, in which tumors of various characteristics invade and metastasize in clusters, has recently been proposed as a poor prognostic factor [[1], [2], [3]]. During migration through the bloodstream as circulating tumor cell (CTC) clusters, cancer cell populations with diverse genetic backgrounds and phenotypes can contribute to the spread of metastasis by enhancing survival from physical stresses, such as blood flow and immune mechanisms [4,5]. Thus, an understanding of the mechanisms governing tumor metastasis is urgently needed to develop effective metastasis-targeted treatment. In this context, based on the established multistep carcinogenesis theory in colorectal cancer, mouse intestinal organoids with various gene mutation combinations were established to conduct animal experiments demonstrating that tumor cells with specific genetic backgrounds have metastatic ability [7]. For example, mouse intestinal tumor cells that have *Apc*^*Δ716*^ (A) and *Trp53*^*R270H*^ (P) gene mutations (abbreviated as AP) showed nonmetastatic, whereas those with *Apc*^*Δ716*^ (A), *Kras*^*G12D*^ (K), *Tgfbr2*^*−/−*^ (T), or *Trp53*^*R270H*^ (P) gene mutations (abbreviated as AKTP) are highly metastatic [6,7]. Moreover, only a specific combination of metastatic and nonmetastatic tumors formed polyclonal metastasis through fibrosis involving sinusoidal vessels due to the interaction between metastatic tumors and hepatic stellate cells (HSCs). However, the detailed dynamics of how the extravascular invasion of polyclonal CTC clusters leads to metastasis formation through metastatic tumor–vessel interactions are still unknown and difficult to clarify in animal experiments [8].

Microphysiological systems (MPS), in which cells are integrated and cultured in arbitrary three-dimensional (3D) shapes on microfabricated devices, are promising experimental tools. MPSs enable the understanding of complex cell–tissue behaviors in the complex cancer microenvironment through a bottom-up constitutive approach. Thus, MPSs may yield information on metastatic mechanisms by mimicking the tumor microvessel (MV) microenvironment to obtain spatiotemporal cell dynamics information from the view point of cell–cell interactions [[9], [10], [11]]. Although several studies have simulated the extravascular invasion process by introducing single cancer cells [[12], [13], [14], [15], [16]], few have investigated this using tumor clusters.

To address the limitations of in vivo studies, we developed a coculture system composed of a 3D MV model with CTC cluster-like tumor organoids inside the MV lumen. We then performed time-lapse imaging to understand the spatiotemporal dynamics of polyclonal extravascular invasion by nonmetastatic AP and metastatic AKTP and the respective contributions of AP and AKTP to this process. We introduced nonmetastatic, metastatic, and mixed organoids into the vasculature and compared their extravascular invasion kinetics.

## 2. Materials and methods

### Cell cultures

#### Endothelial cells

Primary human umbilical vein endothelial cells (HUVECs; Lonza, Basel, Switzerland) were cultured in Endothelial Cell Growth Medium-2 BulletKit (EGM-2; Lonza) at 37°C in a humidified atmosphere with 5% CO_2_. The cells between passages 3 and 4 were used.

#### 2) Hepatic stellate cells

Mouse immortalized Venus-labeled HSCs were obtained from *Trp53* mutant mice, as described previously [7]. HSCs were cultured in high-glucose Dulbecco’s Modified Eagle Medium (DMEM) containing 10% fetal bovine serum (FBS) at 37°C in a humidified atmosphere with 5% CO_2_.

#### 3) Intestinal tumor-derived cells

Mouse intestinal tumor-derived cells were used as previously described [6,7]. Briefly, AP and AKTP cell lines were developed from *Apc*^*Δ716*^ *Trp53*^*R270H*^ and *Apc*^*Δ716*^ *Kras*^*G12D*^ *Tgfbr2*^*−/−*^ *Trp53*^*R270H*^ mouse intestinal tumors, respectively. AP and AKTP cells were labeled with fluorescent proteins tdTomato (AP-tdt, AKTP-tdt) and Venus (AKTP-v), respectively. The tdTomato and Venus cDNAs were subcloned into the pPB-CAG-IP PiggyBac transposon expression vector and cotransfected with the transposase expression vector into cells using Lipofectamine (Thermo Fisher Scientific, Waltham, MA, USA). Consequently, transfected clones were selected by drug selection with 1 μg/mL puromycin (InvivoGen, San Diego, CA, USA). *Mycoplasma* testing was performed using an indirect immunofluorescence test. These tumor cells were cultured on dishes with tumor 2D medium (Advanced DMEM/F-12 medium; Gibco, Thermo Fisher Scientific) supplemented with 10% FBS, 5% penicillin–streptomycin solution (Fujifilm Wako, Osaka, Japan), 5 μM ALK inhibitor (A83-01, Sigma-Aldrich, Saint Louis, MO, USA), 5 μM GSK3 inhibitor (CHIR99021, Sigma-Aldrich), and 10 μM ROCK inhibitor (Y27632, Fujifilm Wako). The cells between passages 3 and 20 were used.

All animal experiments were approved (approval number 2705) by the Committee on Animal Experimentation of Kanazawa University.

### Formation of tumor organoids

Tumor cells were suspended in the 2D medium described above at 5.0 × 10^3^ cells/mL and then seeded into ultralow attachment 384-well round-bottom plates (Sumitomo Bakelite, Tokyo, Japan) at 100 cells/well to form organoids. For the culture of AP/AKTP mosaic organoids, AP and AKTP suspensions were mixed at a 1:1 ratio. The cells were cultured for approximately 60 h, collected in a 1.5-mL tube, and centrifuged three times at 6000 rpm. The supernatant was carefully removed, and the organoids were suspended in fresh media for further experiments.

### MV model

In-house polydimethylsiloxane-based chips (25 mm × 25 mm × 5 mm) were used to prepare the 3D MV model, as described previously [17,18]. A bovine serum albumin (BSA)-coated acupuncture needle (200 μm in diameter; Seirin, Shizuoka, Japan) was inserted into the chip to form a lumen structure in the type I collagen gel (2.4 mg/mL, Cellmatrix type I-A, Nitta Gelatin, Osaka, Japan).

The chips were incubated under a humidified atmosphere with 5% CO_2_ at 37°C for 60 min. HUVECs were seeded into the collagen gel channel with media containing 3% dextran (*Leuconostoc* spp., Mr 450,000–650,000, Sigma-Aldrich) at a density of 1 × 10^7^ cells/mL. After 10 min, the media were replaced with EGM-2 and changed every 1–2 days.

HSC-cocultured 3D MV models were fabricated, as described previously [19]. Briefly, a HUVEC and HSC mixture suspension (1 × 10^7^ cells/mL in total at a 5:1 HUVECs–HSCs ratio) was loaded into the collagen gel channels. The subsequent procedure was the same as described above [19].

### Seeding tumor organoids into an MV model

AP, AKTP, or AP/AKTP organoids suspended in fresh medium were seeded into the MV or HSC-MV lumen from one inlet of the chip. MV or HSC-MV were cultured for up to 7 days. Soon after seeding, the other inlet was closed by the insertion of a BSA-coated acupuncture needle to stop the medium flow. After a few minutes, the settling of organoids in the lumen were confirmed, and the needle was carefully removed. Media were changed every 1–2 days.

### RNA isolation and quantitative RT-PCR

AP and AKTP were cultured on culture dishes as 2D cultures or on ultralow attachment 96-well round-bottom plates (Sumitomo Bakelite) to form 3D organoids. Total RNA was extracted from these cells using the ISOGEN reagent (Nippon Gene, Toyama, Japan), reverse-transcribed using a PrimeScript RT reagent Kit (Takara Bio Inc., Otsu, Japan), and amplified using ExTaqII SYBR Premix (Takara Bio Inc.) on an Aria Mx real-time thermocycler (Agilent Technologies, Santa Clara, CA, USA). The relative mRNA levels of *Adam10, Adam17*, and *Adam28* to the mean *Gapdh* were calculated, and the AKTP values for the figure were compared with those of AP (relative expression = 1). The primer sequences are listed as follows. *Adam10* forward, 5’-TTATGCCATGTTTGCTGCATGA-3’; *Adam10* reverse, 5’-ACTGCTTGCTCCACTGCAAAGA-3’; *Adam17* forward, 5’-ACCTGAACAACGACACCTGCTG-3’; *Adam17* revers, 5’-TTCTGCGCCGTCTCAAACTG-3’; *Adam28* forward, 5’-CTTGAGATGGCCAATTACATCAACA-3’; *Adam28* reverse, 5’-CGGTCCAGATTTCCACTCCAA-3’.

### Immunocytochemistry

Tumor organoids cultured for approximately 60 h were collected in a 1.5-mL tube and centrifuged three times at 6000 rpm. The supernatant was carefully removed, and the organoids were encapsulated in immunoprecipitation gel (GenoStaff, Tokyo, Japan). The organoids were then fixed with 4% paraformaldehyde (PFA) in PBS for 45 min at room temperature, embedded in paraffin, and sliced into 4*-*μm-thick sections for immunofluorescence staining. For immunohistochemistry, antibodies against green fluorescent protein (GFP; #2956, Cell Signaling Technology, Danvers, MA, USA), red fluorescent protein (RFP; #600-401-379, Rockland Immunochemicals Inc., Limerick, PA, USA), E-cadherin (#AF748, R&D, Minneapolis, NE, USA), and Ki67 (#16667, Abcam, Waltham, MA, USA) were used as primary antibodies. GFP and RFP antibodies were used to detect Venus-or tdTomato-labeled cells, respectively. Alexa Fluor 594-or Alexa Fluor 488-conjugated antibodies (1:200, Thermo Fisher Scientific, Waltham, MA, USA) were used as secondary antibodies for fluorescence immunohistochemistry. 4′,6-diamidino-2-phenylindole was used for nuclear staining.

### Microscopy

To observe tumor organoids in MVs, bright-field and fluorescent images were captured using a fluorescenCE microscope (Axio Observer Z1, Carl Zeiss, Oberkochen, Germany) equipped with a 10× or 20× observation lenses. For time-lapse observations, Z-stack images were CAPTURED using a confocal laser scanning microscope (CLSM; LSM700, Carl Zeiss) equipped with a 20× objective lens. To detect cellular dynamics sequentially, Z-stack images were captured using a CLSM at 30-min or 1-h intervals. Red and green fluorescence was detected using laser wavelengths of 488 and 405 nm, respectively. The acquired Z-stack images were exported using IMARIS software (version 9.0.0, BitPlane, Zurich, Switzerland).

### Image analyses

#### Quantification of tumor organoid growth

Z1 images of tumor organoids cultured on ultralow attachment 384-well round-bottom plates for days 1, 2, 3, and 7 were used. The outline of each organoid was detected manually, and the area and perimeter were automatically measured using the Fiji/ImageJ software (ver. 2.90). The average diameter and circularity of each organoid were calculated using the formulas 2(area/π)^1/2^ and 4π(area/perimeter^2^), respectively.

#### Quantification of tumor organoid invasions

The effects of HSCs on tumor invasion into collagen were evaluated using Z-stack images of the AP/AKTP invasion area at day 6 after tumor organoids were seeded into the lumen of MV or HSC-MV. 3D images were reconstructed from Z-stack images and binarized, and the number of specific invasion areas (microfoci) was manually counted using IMARIS software (version 9.0.0). Subsequently, to count spikes, binarized images were displayed with triangle meshes by calculating mean curvature using MeshLab software (ver. 2020.12). The front view of each 3D structure was captured and imported to Fiji/ImageJ, and the number of spikes was manually counted using the Cell Counter Fiji software plugin. Microfoci and spikes were counted at three independent captured areas of one sample in each culture condition, and the total numbers in each culture condition were then divided by the total captured length of MV/HSC-MV.

### Statistical analysis

The qPCR data were analyzed using two-tailed unpaired Student’s *t*-test using Graphpad Prism 10.0.2 (Boston, MA, USA). Values are presented as mean ± standard deviation (s.d.). Differences between means were considered statistically significant at *p* < 0.05. All data were reproduced by performing at least two independent experiments.

## 3. Results

### 3-1. Development of an extravasation tumor–MV model

To investigate the mechanism of polyclonal metastasis, an in vitro extravasation tumor–MV model was developed. Nonmetastatic AP, metastatic AKTP, and AP/AKTP mosaic tumor organoids mimicking CTC clusters were prepared by seeding into ultralow attachment round-bottom wells at the same cell density. The cells aggregated within 24 h, and the size and circularity increased and became stable over time, respectively (Fig. 1A and 1B). Interestingly, the mosaic organoids showed an intermediate phenotype between AP and AKTP. The AKTP organoids showed a lower aggregation tendency than AP organoids, resulting in a larger size and nonuniform structure during the initial culture period. Histological analyses of the organoids on day 3 showed that AP, AKTP, and AP/AKTP organoids maintained structures with cell–cell junctions expressing E-cadherin. However, more Ki67-positive proliferative cells were observed in the organoids containing AKTP cells (Fig. 1C). The tumor organoids were introduced into the MVs on day 3 to investigate further interactions between the tumors and MVs under microscopy (Fig. 1D).

**Fig. 1.**
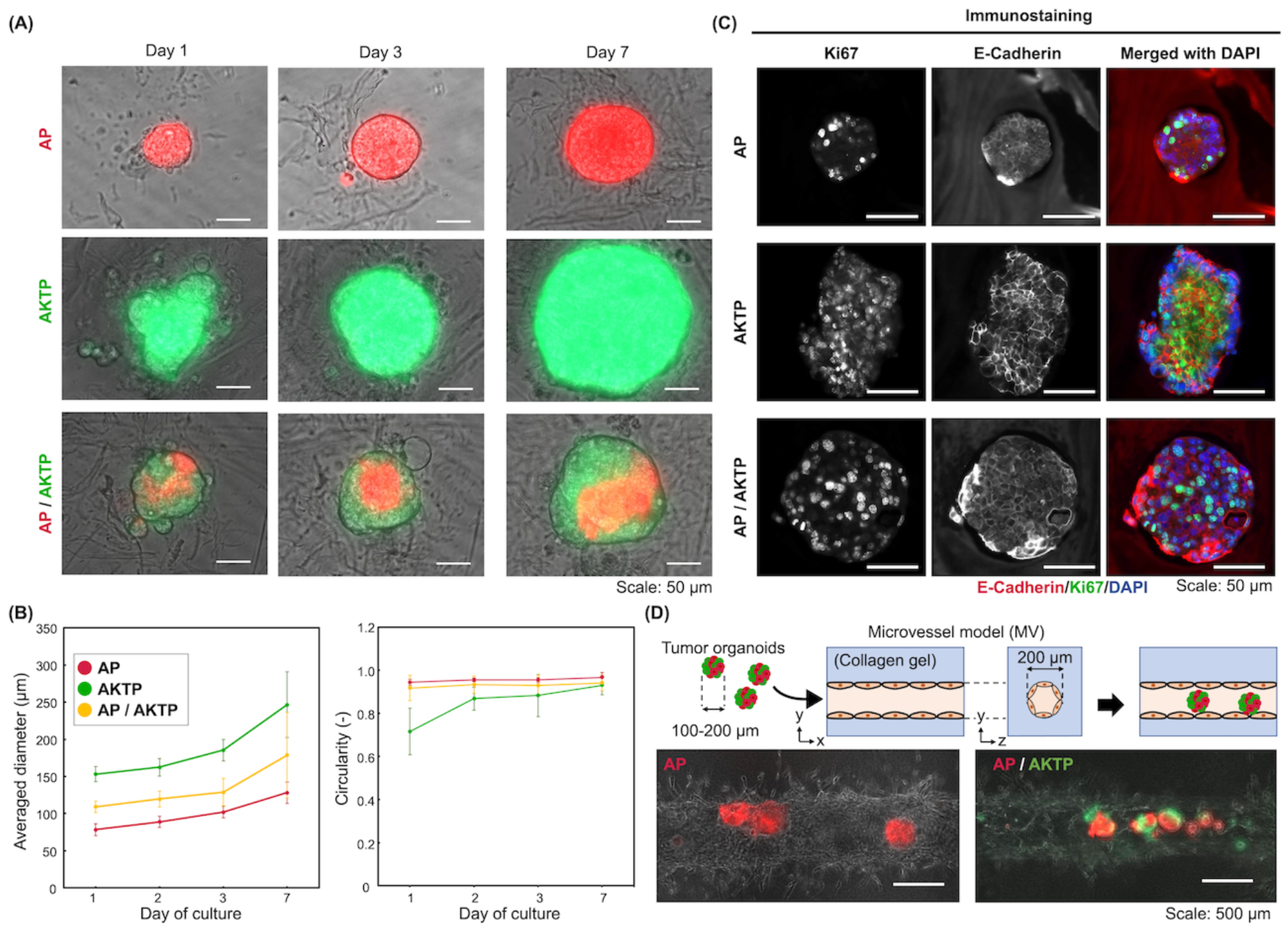
Development of the extravasation tumor–MV model. **(A)** Time-dependent microscopic views of tumor organoids cultured in ultralow attachment plates. **(B)** Time-dependent diameter and circularity of organoids. Mean ± standard deviation (n = 3). **(C)** Immunofluorescent images of tissue sections of tumor organoids on day 3. **(D)** Microscopic images of MVs the day after introducing tumor organoids into the lumen.

### 3-2. Role of metastatic tumors during polyclonal metastasis for extravasation

Noninvasive AP, invasive AKTP, or AP/AKTP organoids introduced into the MV showed sequential invasive steps of organoids over the culture period: (i) vessel co-option, (ii) vessel constriction or destruction, and (iii) tumor extravascular invasion (Fig. 2A). The organoids containing metastatic AKTP clearly exhibited more invasive behavior. Nonmetastatic AP organoids hijacked along the MV wall (Fig. 2A, top left) resulted in vessel co-option, even after 28 days in culture (data not shown). In contrast, metastatic AKTP showed vessel co-option but remained filled with divided small clusters in the luminal space on day 14. Furthermore, the MV lumen was locally constricted on day 21 (Fig. 2A, top right). Similar to the AKTP organoids, the AP/AKTP mosaic organoids likewise invaded the inner walls of MVs during the initial culture period. Constrictions with micrometastasis-like aggregates of both AP and AKTP were locally observed on day 14 (Fig. 2A, bottom). Finally, on day 21, the constriction domains expanded, and extravascular invasion into the outer collagen gel with tumor spike formation was observed (Fig. 2A, bottom).

**Fig. 2.**
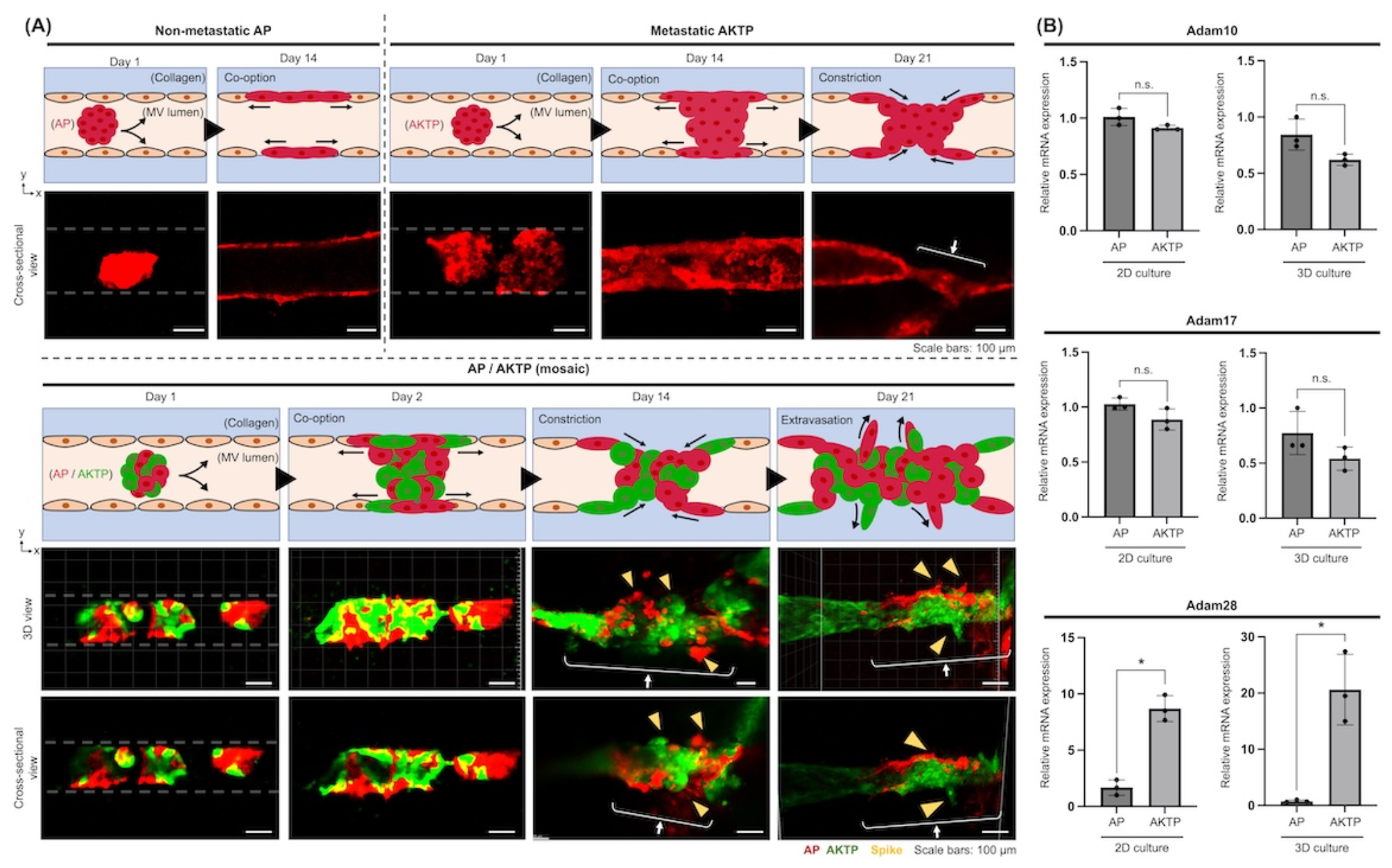
Spatiotemporal observation of the tumor organoid in microvessels (MVs). **(A)** Time-lapse images of organoid invasion obtained by confocal laser scanning microscopy: (left) nonmetastatic AP, (right) metastatic AKTP, and (bottom) AP/AKTP in MVs. White dotted line: MV layers; white arrows: aggregation of tumor cells through MV lumen constriction; yellow arrowheads: spikes of tumor cells. **(B)** Relative mRNA levels of disintegrin and metalloproteinase family enzymes in 2D or 3D cultures of nonmetastatic AP and metastatic AKTP (n = 3). Mean ± standard deviation. \**p* < 0.05. n.s., no significant difference.

To investigate the molecular mechanism of vascular remodeling induced by metastatic AKTP, we analyzed gene expression of the a disintegrin and metalloproteinase (ADAM) protein family member related to extracellular matrix (ECM) degrading enzymes. Quantitative reverse-transcription polymerase chain reaction showed that *Adam28* expression was significantly higher in metastatic AKTP in both 2D and 3D cultures (Fig. 2B).

### 3-3. Dynamics of metastatic tumor HSCs during extravasation

Our previous in vivo studies demonstrated that metastatic AKTP secretes transforming growth factor-β1 to induce HSCs to form malignant fibrotic niche, thereby promoting colonization into liver tissue, whereas nonmetastatic AP could not and subsequently failed to induce metastasis in the liver [7]. To understand the changes in the extravascular fibrotic niche induced by tumor organoids during extravasation, the spatiotemporal dynamics between AP or AKTP tumor organoids and HSCs were investigated using the HSC-MV model. A mixed cell suspension of mouse HSCs and HUVECs introduced into the collagen gel luminal structure successfully formed endothelium layers surrounded by HSCs (Fig. 3A).

**Fig. 3.**
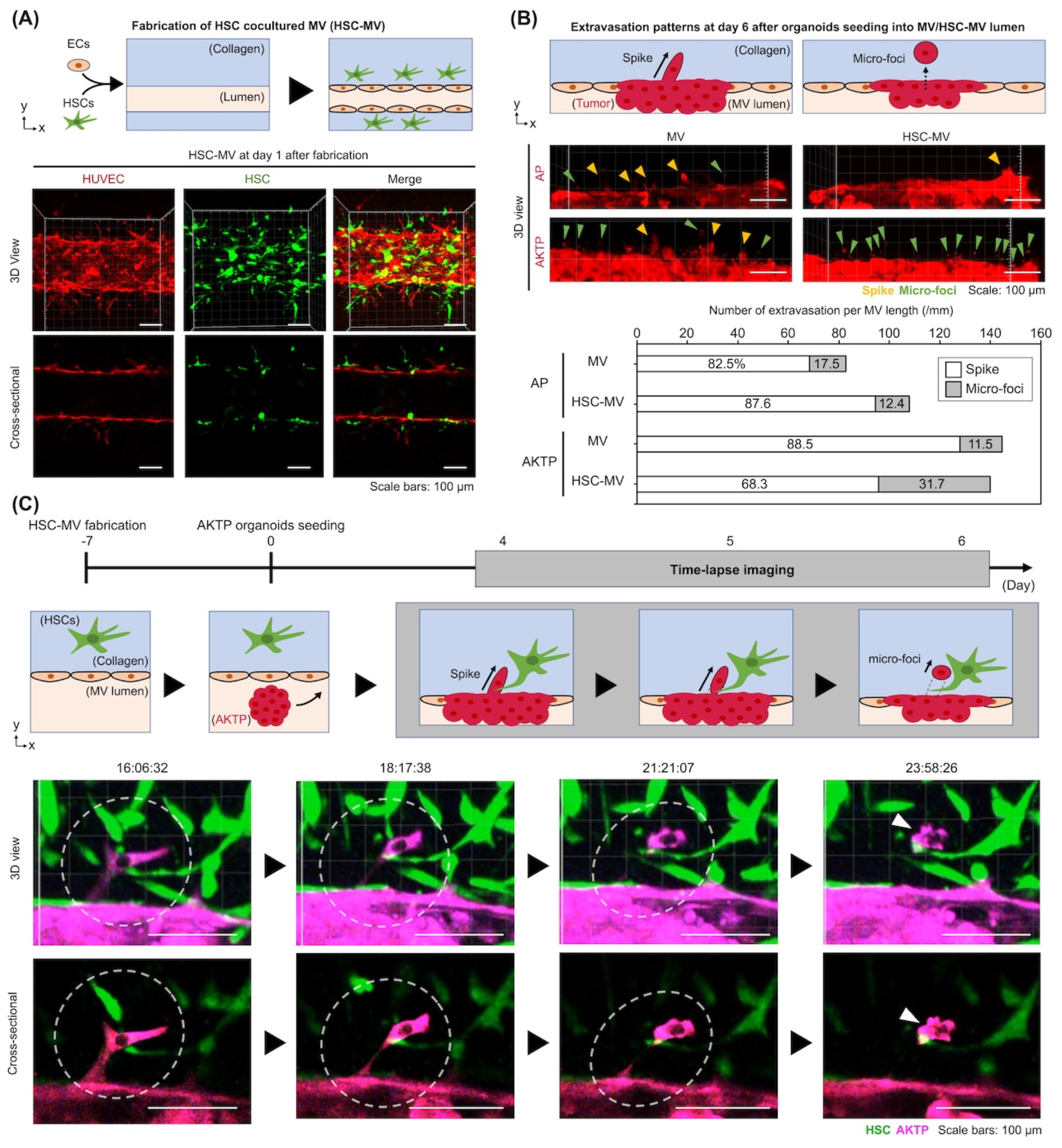
Spatiotemporal observation of tumor organoids in hepatic stellate cells–microvessel models (HSC-MVs). **(A)** Confocal laser scanning microscopy (CLSM) images of HSC-MVs on day 1 after fabrication. HSCs localized around the MVs. **(B)** Extravasation patterns of tumor organoids from MVs or HSC-MVs. Schematic illustration of the extravasation pattern (top). CLSM images of extravasation of tumor cells from co-option in MV or HSC-MV on day 6 (middle). Yellow arrowhead: extravascular tumor spike; green arrowhead: tumor microfoci. Number of spikes and microfoci of nonmetastatic AP and metastatic AKTP in MV and HSC-MV at day 6 (bottom). **(C)** Time-lapse imaging of the extravasation of metastatic AKTP inducing microfoci formation via interaction with HSCs. Schematics of metastatic AKTP organoids seeding into the HSC-MV lumen for time-lapse imaging (top); and representative time-lapse images of metastatic AKTP invasion into the collagen gel area in HSC-MV at days 4–6 (bottom). White dotted circles: extravascular spike; white arrowhead: tumor microfoci.

AP or AKTP organoids introduced into the MV or HSC-MV exhibited vessel co-option. Extravascular spike and tumor microfoci formations, which consisted of independent cells released from the co-opted area, were observed on day 6 (Fig. 3B). Metastatic AKTP has a tendency to increase tumor microfoci formation (31.7%) in HSC-MV at approximately threefold of that in MV (11.5%). In contrast, for nonmetastatic APs, no trend was observed in the tumor microfoci formation percentage during HSC coculture (12.4%) compared with that without HSCs (17.5%). Furthermore, time-lapse imaging captured the sequential formation of micrometastases (tumor microfoci) from the tip of the spike via direct metastatic AKT–HSC interaction: (i) extravascular spike formation by AKTP cells, (ii) HSCs touching the tip of the spike, and (iii) release of AKTP cells, resulting in tumor microfoci formation (Fig. 3C).

## 4. Discussion

An in vitro extravasation tumor–MV model was developed to understand the mechanism of polyclonal metastasis using tumor organoids with different driver mutations: nonmetastatic AP, metastatic AKTP, and AP/AKTP mosaic organoids. Organoids, including metastatic AKTP, exhibited extravasation of tumors via vessel co-option, constriction, and destruction, whereas nonmetastatic AP exhibited only vessel co-option (Fig. 2A). In particular, the in vitro experimental system revealed for the first time that mosaic organoids maintained their mosaic state during the whole extravasation process, from the introduction into the MV lumen to the formation of colonization-like clusters. These are consistent with the results of a previous study involving mice experiments in which nonmetastatic APs formed polyclonal metastatic nests together with metastatic AKTP [7].

We also found that AKTP cells highly expressed *Adam28*, which is involved in cancer proliferation, invasion, and metastasis and thus confers the ability to alter the vascular microenvironment (Fig. 2B). *Adam28* is a member of the *ADAM* gene family, which is involved in cancer invasion and metastasis through the degradation of von Willebrand factor. These proteins upregulate the active forms of insulin-like growth factor-1, which contributes to cell proliferation and apoptosis evasion [[20], [21], [22], [23]]. In addition, some genetic nucleotide sequences of *Adam28* encode protease domains similar to those of matrix metalloproteinases, which may be crucial to extravascular invasion [24]. Therefore, vascular remodeling induced by metastatic AKTP may involve ECM degradation caused mainly by *Adam28* expression.

In addition, extravascular invasion dynamics observed in HSC-MV models revealed the formation of foci-like micrometastases from the tips of protrusions via a direct interaction between metastatic AKTP and HSCs (Fig. 3C), which could not be clarified in a previous mouse experiment [7]. Furthermore, evaluation of the number of foci-like formations indicated that the number of foci-like formations tended to triple in metastatic AKTP, whereas no increase in foci-like formations was observed in nonmetastatic AP (Fig. 3B). This difference in the interactions of HSCs with foci-like formations between nonmetastatic and metastatic tumors may confirm previously reported results describing the metastatic ability of each tumor cell type [7]. Therefore, a complex heterogeneous environment accelerates microfoci formation. HSCs interact with the tip of the extravascular spike to form microfoci.

## 5. Conclusion

In conclusion, we developed in vitro extravasation tumor-MV model to understand the mechanism of polyclonal metastasis using mouse intestinal tumor-derived organoids with different driver mutations. Time-lapse imaging determined the sequential steps of the extravasation of tumor cells via vascular remodeling. Metastatic tumor facilitated vascular remodeling, resulting in extravascular invasion. The extravasation dynamics by tumor organoids with vessel co-option and spikes have been discussed almost exclusively in the context of intravasation. They differ fundamentally from the transendothelial migration processes involved in the conventional single-cell extravascular invasion mechanism, followed by adhesion of intercellular adhesion molecule-1 to the vessel wall on the vascular endothelial membrane and invasion outside the blood vessels by passing through the vascular endothelial cell junctions [[25], [26], [27]]. The novel extravasation pattern observed in this model may lead to the discovery of novel targets and the design of therapeutic strategies for drug delivery systems targeting metastatic sites [28].

## Author contributions statement

**Yukinori Ikeda:** Writing: original draft, investigation, funding acquisition, Formal analysis, Data curation. **Hiroko Oshima:** Writing – original draft; Investigation; Software; Data curation; Resources. **Kok Sau Yee:** Writing – original draft; Investigation; Software; Data curation; Resources. **Masanobu Oshima:** Writing – review & editing; Writing – original draft; Supervision; Resources; Project administration; Funding acquisition; Data curation; Conceptualization. **Yukiko T Matsunaga:** Writing, review and editing; Writing – original draft; Supervision; Resources; Project administration; Funding acquisition; Data curation; Conceptualization.

## Acknowledgments

This work was supported by AMED P-CREATE (JP21cm0106272); the Extramural Collaborative Research Grant of Cancer Research Institute, Kanazawa University; JSPS KAKENHI Grant-in-Aid for JSPS Fellows (JP23KJ0490); the Sasakawa Scientific Research Grant from The Japan Science Society; and WINGS-QSTEP.

